# Distilling Mechanistic Models From Multi-Omics Data

**DOI:** 10.1101/2023.09.06.556597

**Authors:** Samantha Erwin, Joshua R. Fletcher, Daniel C. Sweeney, Casey M. Theriot, Cristina Lanzas

## Abstract

High-dimensional multi-omics data sets are increasingly accessible and now routinely being generated as part of medical and biological experiments. However, the ability to infer mechanisms of these data remains low due to the abundance of confounding data. The gap between data generation and interpretation highlights the need for strategies to harmonize and distill complex multi-omics data sets into concise, mechanistic descriptions. To this end, a four-step analysis approach for multiomics data is herein demonstrated, comprising: filling missing data and harmonizing data sources, inducing sparsity, developing mechanistic models, and interpretation. This strategy is employed to generate a parsimonious mechanistic model from high-dimensional transcriptomics and metabolomics data collected from a murine model of *Clostridioides difficile* infection. This approach highlighted the role of the Stickland reactor in the production of toxins during infection, in agreement with recent literature. The methodology present here is demonstrated to be feasible for interpreting multi-omics data sets and it, to the authors knowledge, one of the first reports of a successful implementation of such a strategy.

## 1 Introduction

Technological advances have enabled the rapid acquisition of multiple types of omics data from a single biological sample, known as multi-omics. Furthermore, time course collection of these types of samples provides a rich data set that potentially allows an increasingly complete picture of the system dynamics. This data collection strategy can be implemented within a host organism, water, soil, or air. While these approaches are becoming more prevalent, leveraging these rich datasets to understand time course interactions is still limited. Through mathematical analysis, we can better understand early warning markers and metabolic pathways that lead to disease.

To leverage rich multi-omics data sets, network analysis and graphical models have emerged as especially useful frameworks for analyzing the high-dimensional data acquired from multiomics assays by representing these data as a series of discrete, interconnected nodes [1, 2]. Implementing these graphical models enable visualization of the strongest correlations between DNA sequences, transcripts, proteins, metabolites and other data to highlight important relationships or connections which may otherwise be obfuscated by the large quantities of data collected in these assays [3]. Once identified, these connections yield roadmaps which can be used to develop mechanistic models of the biological processes being studied. Despite the increasingly rich landscape of high-dimensional data, the techniques and strategies currently available to maximize biological insights from these data remain underdeveloped. Specifically lagging is the ability to elucidate control mechanisms for infection by developing parsimonious mechanistic models from multiomics data.

One method to mathematically represent causal relationships in complex biological mechanisms is parsimonious ordinary differential equation (ODE) models. These models require relatively little computational overhead and elegantly express biological phenomena in a manner that can be easily manipulated without exhaustive experimentation. For example, ODE models have been used to highlight viral dynamics and infection in host organisms [4], characterize latent infection in viruses [5, 6, 7], T cell death rates [8, 9], and better understand drug therapies for infected patients [10, 11, 12]. However, it is often challenging to accurately determine the connectedness of the ODE systems in biological specimens, given the large number of complex interactions occurring simultaneously.

To address this gap, we have developed a pipeline to identify critical relationships in high dimensional omics data, with the goal of developing mechanistic models of the underlying biological processes. We demonstrate our methodology using a time course model of a *Clostridioides difficile* murine infection [13], characterizing changes in the metabolic landscape and *C. difficile* transcriptome before challenge with *C. difficile* bacteria, during colonization, and after the onset of disease. To this end, we used a sparse penalization method to identify key metabolites in vivo that are strongly associated with toxin production. These relationships were used to develop a mechanistic ODE model of the interaction between nutrients and toxin production. Specifically, a graphical lasso algorithm identified proline and 5-aminovalerate as key metabolites strongly correlated with toxin gene expression. From this, a mechanistic ODE model was developed to describe metabolic and gene expression dynamics (transcriptome) and calibrated using a Markov chain Monte Carlo (MCMC) fitting method to normalized data. Using a sensitivity analysis, we identify novel proline pathways mechanistically that affect toxin expression and the course of infection. This study provides a novel pipeline to develop and parameterize a parsimonious mechanistic model using large-scale omics data and serves as a first step to use metabolomic data in a mechanistic model. Future work is needed to apply this type of methodology to a variety of other data sets to show the robustness of the approach.

## 2 Methods

Our proposed method has four main analysis steps for combining multi-omics data into an interpretable mechanistic model (Figure 1). Each are described in detail below, but are summarized here:

1. Data harmonization
2. Induce sparsity
3. Develop mechanistic models
4. Interpretation

**Figure 1.**
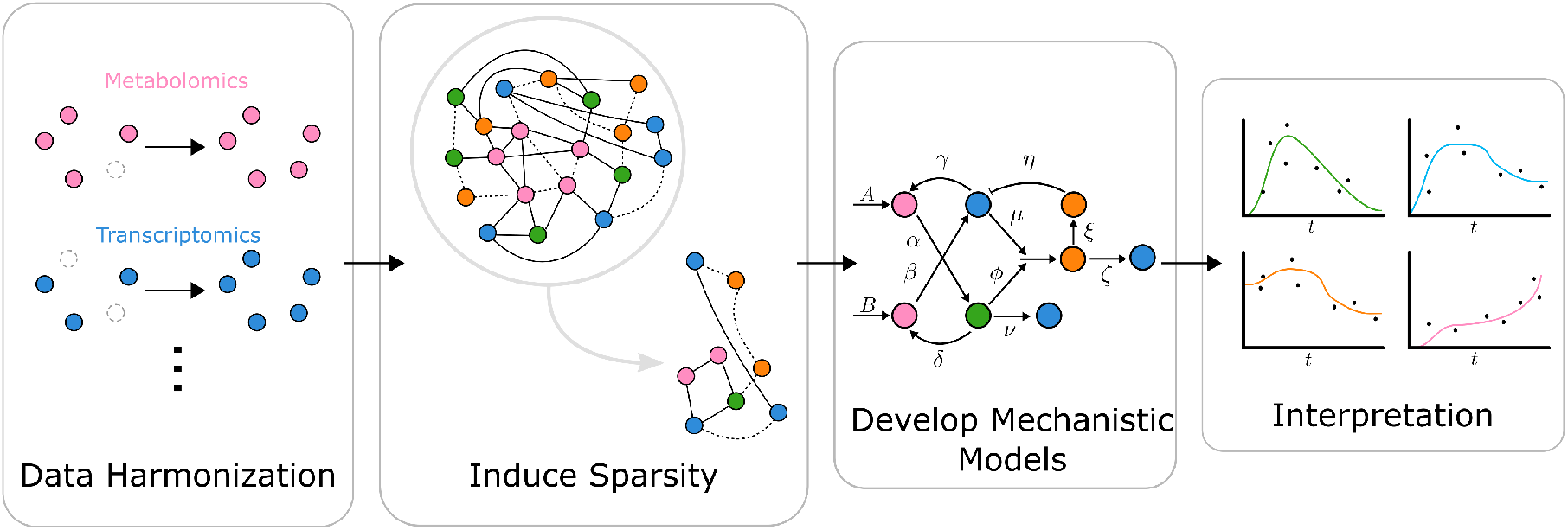
Proposed methodological pipeline

### 2.1 Data harmonzation through metabolomics manipulation

Metabolomic and transcriptomic data from an acute mouse model of *C. difficile* infection was used as the basis for the present study[13]. In this work, 32 mice were initially treated with the antibiotic cefoperazone in their drinking water to render them susceptible to *C. difficile* infection. The mice were then challenged with *C. difficile* VPI 10463 spores and euthanized at different times post-challenge (0, 12, 24, and 30 h); eight mice for each time point. Cecal content was processed using global metabolomic analysis (Metabolon, Inc, Durham, NC). The relative abundance of 638 metabolites were measured including lipids, xenobiotics, carbohydrates, nucleotide, amino acid, cofactors and vitamins, energy, and peptides. Additionally, toxin gene expression levels were determined via quantitative reverse transcription PCR (qRT-PCR) using primers specific to the C. difficile toxins, *tcdA* and *tcdB* genes, as well as the housekeeping gene *rpoC* for normalization. Additional information on the experimental methods, including the raw data, can be found in [13].

Historically, metabolomics data have been reported in relative abundance, where raw data are measured in terms of ion counts for each unique metabolite which are normalized across all samples, scaling the median value to one. The raw metabolomics data was normalized based on ion counts in the present work. Missing values are inferred with the minimum values of the specific metabolite across all samples, which is a standard industrial practice, under the assumption that all missing values are due to an issue with the detection limit rather than a sample handling issue [14, 15, 16]. In the present study, 251 of 638 unique metabolites had missing data for at least one sample. A total of 1976 metabolite measurements were missing from the total 20416 metabolite measurements, or 9.7% across all samples.

More recently, it has been hypothesized that missing metabolomics data are more likely the result of sample processing issues during collection and analysis, rather than issues with instrument sensitivity [17]. Taking this into consideration, we used a K-nearest neighbor analysis to impute the missing data [18]. This method assumes the missing value will take on values similar to neighboring samples, rather than assuming a sample is near the instrument detection limit. Imputation techniques were previously compared using data processed by Metabolon, Inc. and the K-nearest neighbor algorithm was demonstrated to be the most robust approach to reconstruct metabolic pathways and increase statistical power [18]. Additional methods for imputing missing values are available, though it is generally necessary to address missing values prior to sample analysis [19]. Therefore, we combined the metabolomic data with the *in vivo tcdA* and *tcdB* transcriptomic dataand filled in missing data using the K-nearest neighbor algorithm.

### 2.2 Induce sparsity with sparse graphical models

Metabolomics data are often high-dimensional, as the number of measured variables exceeds the number of samples. High dimensionality introduces analysis and interpretation challenges, including accumulation of noise and spurious associations [20]. Sparse graphical models offer an intuitive way to represent dependencies across variables in large data sets with nodes representing variables and the edges connecting the nodes indicating the dependencies between the variables (e.g. pairwise correlations). Sparsifying these graphs reduces the number of interactions to its most highly interdependent components [21, 22, 23]. Furthermore, the process of sparsifying high dimensional data removes unrelated or insensitive interactions, enabling the reduction of these data to more practically useful, identifiable models. Numerous methodologies to sparsify omics data have been proposed, including the sparse partial least-squares discriminant analysis [24], which we previously used to combine metabolomic and transcriptomic data [13].

Arguably, the most popular methodology to induce sparsity into high-dimensional data involves using a penalized log-likelihood function and has been adapted into numerous forms and computational packages [25]. The penalized log-likelihood function is based on the least absolute shrinkage and selection operator (lasso) which minimizes the residual sum of squares based on the sum of the absolute values of the coefficients being less than the selected constant, *λ* [26]. This *L*_1_ penalty method is then used to estimate sparse undirected graphical models by fitting a lasso model [27]. The traditional graphical lasso (glasso) estimates a sparse inverse covariance matrix, Θ, using an *L*_1_ penalty, *λ*, and the sample covariance matrix, *G*. Thus, it minimizes the log-likelihood

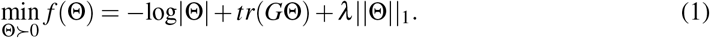

In the present work, the glasso was implemented using a correlation matrix for **G** [28]. We calculate the correlation matrix using Spearman’s rank, which measures the degree of association between ranked random variables. Using **G** gives us the sparse inverse correlation matrix, Θ, which is transformed into the penalized partial correlation matrix, **R**, and only contains the strongest partial correlations. The entries in Θ, *θ*_*i j*_, are used to calculate the penalized partial correlation matrix, **R**,

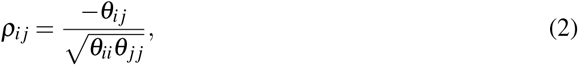

where *ρ*_*i j*_ are entries in **R**. Using, **R**, the iGraph package was used to develop a graphical model in which each variable is represented as a node (i.e. lipid, carbohydrate, xenobiotics, nucleotide, amino acid, energy, peptide, cofactors, and toxin) interconnected by edges weighted by the partial correlations between variables [29]. This graph was then used to guide the development of mechanistic ODE models of the interactions between metabolites and toxin production.

### 2.3 Mechanistic model identifiability and parameter estimation

Following the sparsification process **G** *→* **R** described in Section 2.2, **R** contains only the partial correlations above the threshold set by *λ* . This process is analogous to other re-parameterization techniques which greatly reduce the parameter space from which to glean mechanistic information [30]. In conjunction with this parameter space reduction, we leveraged *a priori* biological knowledge about possible related parameters, including the causal relationship between toxin production and the pathogenesis of *C. difficile* colonization [31], evidence that *C. difficile* colonization is correlated to specific metabolites [32] and relies on electron donors for energy [33]. Together, *a priori* knowledge and identifying the strongest relationships between metabolites and transcripts were used to inform the practical identifiability of a schematic representation of metabolite-mediated toxin production during *C. difficile* colonization (Figure 4).

From this schematic representation, we developed a mechanistic ODE model with nodes that remain highly connected to the disease state of interest chosen as variables. Writing the ODE model representing these connections in a normalized fashion simplifies relationship determination between data measured in concentration and improves numerical stability, similarly to other studies [34]. The ODE model was implemented using an LSODA-based solver [35] andfit to the selected metabolite concentrations at each sample time using an affine-invariant MCMC algorithm [36].

*A posteriori* analysis reinforced the identifiability of the ODE model from the normalized metabolomic and transcriptomic data. First, a sensitivity analysis was performed on the ODE system using a combination of Latin hypercube sampling and partial rank correlation coefficients (PRCC) [37]. These analysis identifies the parameters whose variation has the most influence over the model dynamics (Figure A8) and are discussed in greater detail in Appendix A. Further evidence for the practical identifiability of the ODE model is provided by the convergence of MCMC chains which is discussed in Section 3.2.2.

### 2.4 Model analysis and interpretation

Following model development and parameter estimation, the model can then be used to understand the specific metabolomic mechanisms driving disease, namely *C. difficile* infection, in the present case. By parameter variation, a sensitivity analysis is performed to better understand the specific mechanisms leading to the target of interest [38] and identify potential therapeutic targets. All statistical analyses were performed using R version 3.4.3 and R studio version 1.2.1335 [39, 40] while the ordinary differential equation models were solved numerically using Scipy v1.10.1 with parameter fitting performed using LMFIT v1.2.1 in Python v3.10.6 [41].

## 3 Results

### 3.1 Sparse graphical models of interactions between toxins and metabolites

The glasso algorithm with *λ* = 0.8 was used to sparsify the combined data set (Figure 2). This threshold was selected after characterizing how the total number of metabolites and total number of metabolites specifically connected to *tcdA* or *tcdB* varied as the *λ* was increased to balance the sparsity and information retention. The partial correlations between the expression levels of toxin genes *tcdA* and *tcdB* and metabolites when *λ* = 0.8 are provided in Table 1. It is important to emphasize that the *λ* parameter was selected to remove weakly-correlated elements from the graph. 5-aminovalerate and proline were highly correlated with both *tcdA* and *tcdB*, in agreement with previous results based on correlations between time points [13]. This also corroborates work that has suggested that the Stickland reaction is necessary for robust *C. difficile* growth [42, 33, 43, 44]. Hydroxyphenyl propionate, guanosine, and sedoheptulose were also partially correlated with *tcdA* and *tcdB*.

**Table 1:** Using the penalty *λ* = 0.8 we select the subset of metabolites connected to *tcdA* and *tcdB*. We list the weights of the partial correlation from the metabolite to the specific toxin in the column next to the specific metabolite. Five consistent metabolites were found to be connected or partially correlated to *tcdA* and *tcdB* at all time points during the study: 2-4-(hydroxyphenyl) propionate, 5-aminovalerate, guanosine, proline, sedoheptulose.

| *tcdA* connection | weight | *tcdB* connection | weight |
| --- | --- | --- | --- |
| 2-4-(hydroxyphenyl) propionate | -0.03502 | 2-4-(hydroxyphenyl) propionate | -0.03394 |
| Hydroxybutyrate | 0.01046 | 5-aminovalerate | 0.034087 |
| 5-aminovalerate | 0.02467 | Guanosine | -0.00679 |
| Equol sulfate | 0.00245 | N-acetylleucine | -0.00417 |
| Guanosine | -0.00736 | Proline | -0.01658 |
| Proline | -0.01046 | Sedoheptulose | -0.01309 |
| Sedoheptulose | -0.02311 |  |  |

**Figure 2.**
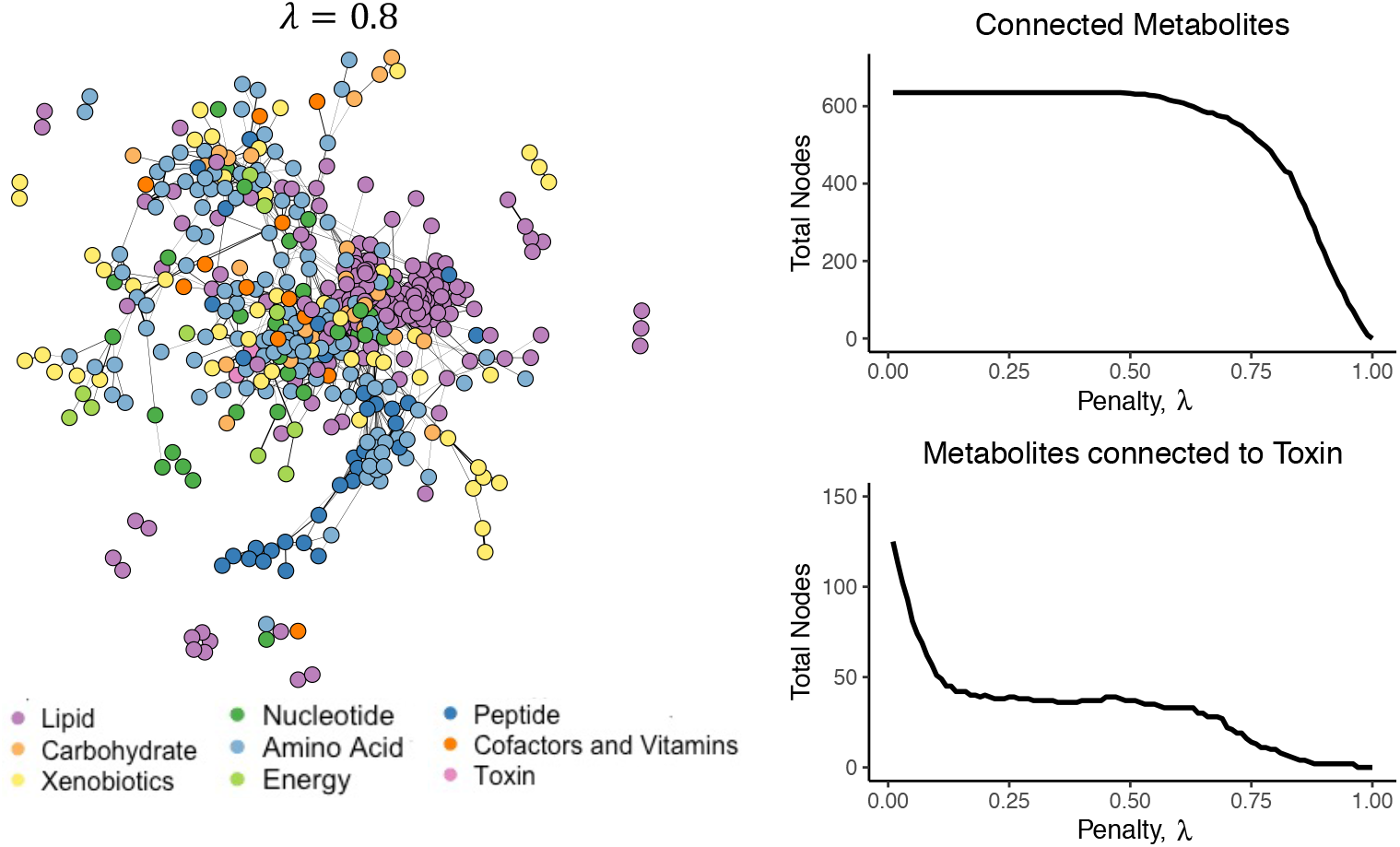
Graph using metabolomic data measured in raw ion counts with missing values filled in using the K-nearest neighbor algorithm. The left panel shows a sparse graph with *λ* = 0.8. The graphs in the right panel show the total nodes are color-coded across all graphs based on the subclasses indicated in the legend. The solid lines indicate a positive partial correlation and the dashed lines a negative partial correlation. Thicker lines indicate a stronger partial correlation. The right panel shows a graph of the total metabolites in a graph as *λ* is increased (top) and the metabolites connected to toxin as *λ* is increased (bottom)

A graphical representation of the metabolites with high partial corrrelations to *tcdA* and *tcdB* expression listed in Table 1 are shown in Figure 3 with solid lines indicating positive correlations, dashed lines indicating negative correlations, and line thickness representing the strength of the correlation. These graphical results visually highlight the negative partial correlation of hydroxyphenyl propionate, guanosine, and sedoheptulose. We also see strong negative partial correlations with proline indicating that as proline concentration decreases, *tcdA* and *tcdB* expression increases. Furthermore we see positive partial correlations between 5-aminovalerate and *tcdA* and *tcdB* which indicates that as their expression increases, 5-aminovalerate concentration also increases. This is consistent with the known biology of C. difficile, which ferments proline to 5-aminovalerate as a preferred mechanism of ATP generation [45, 46, 47, 48, 49, 50, 51].

**Figure 3.**
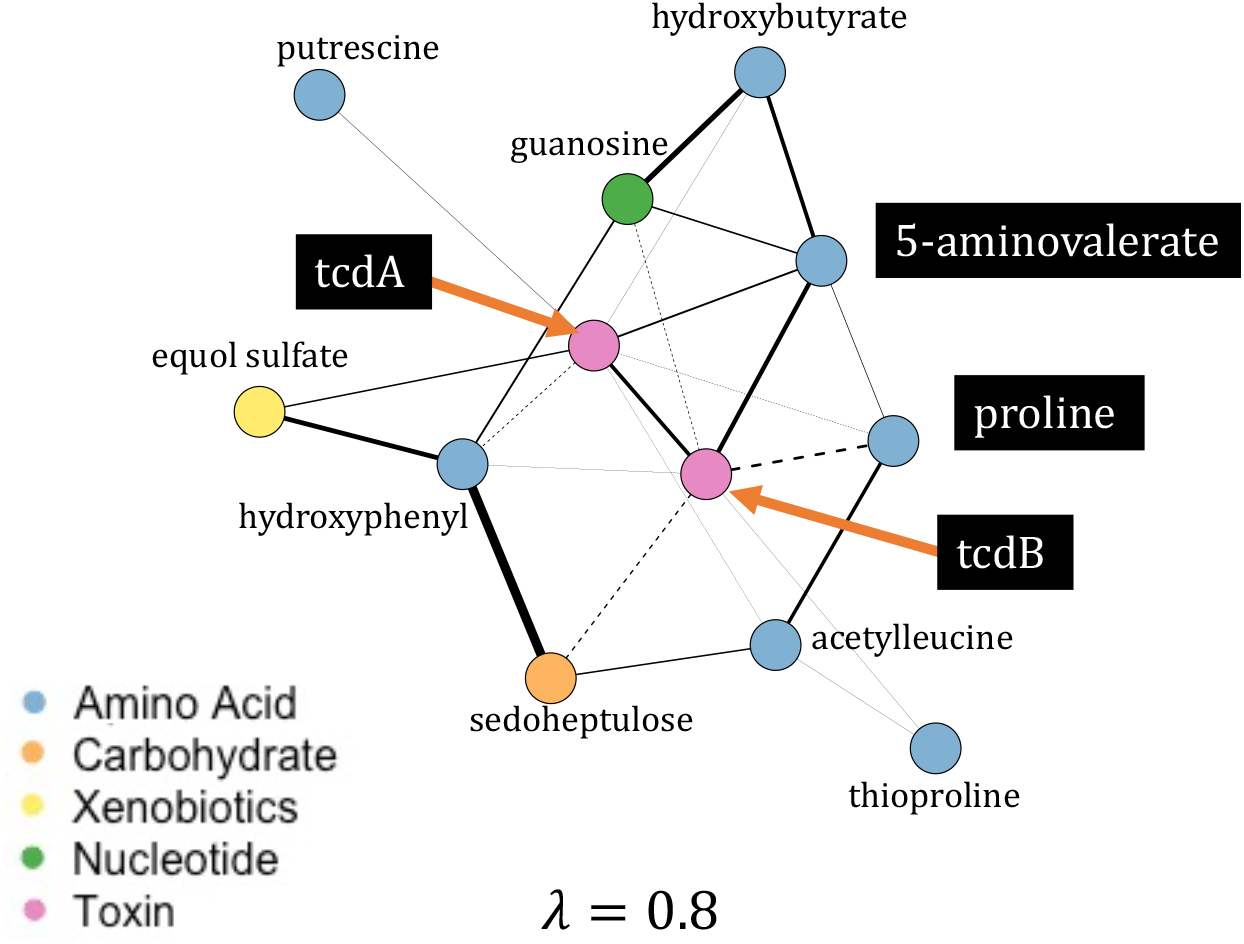
Specific metabolites connected to toxin in a sparse sub-graph with a penalty of *λ* = 0.8. The nodes are color-coded based on subclass indicated in the legend. The solid lines indicate a positive partial correlation and the dashed lines a negative partial correlation. Thicker lines indicate stronger partial correlations. The metabolites names are listed, with an emphasis on *tcdA, tcdB*, 5-aminovalerate and proline.

**Figure 4.**
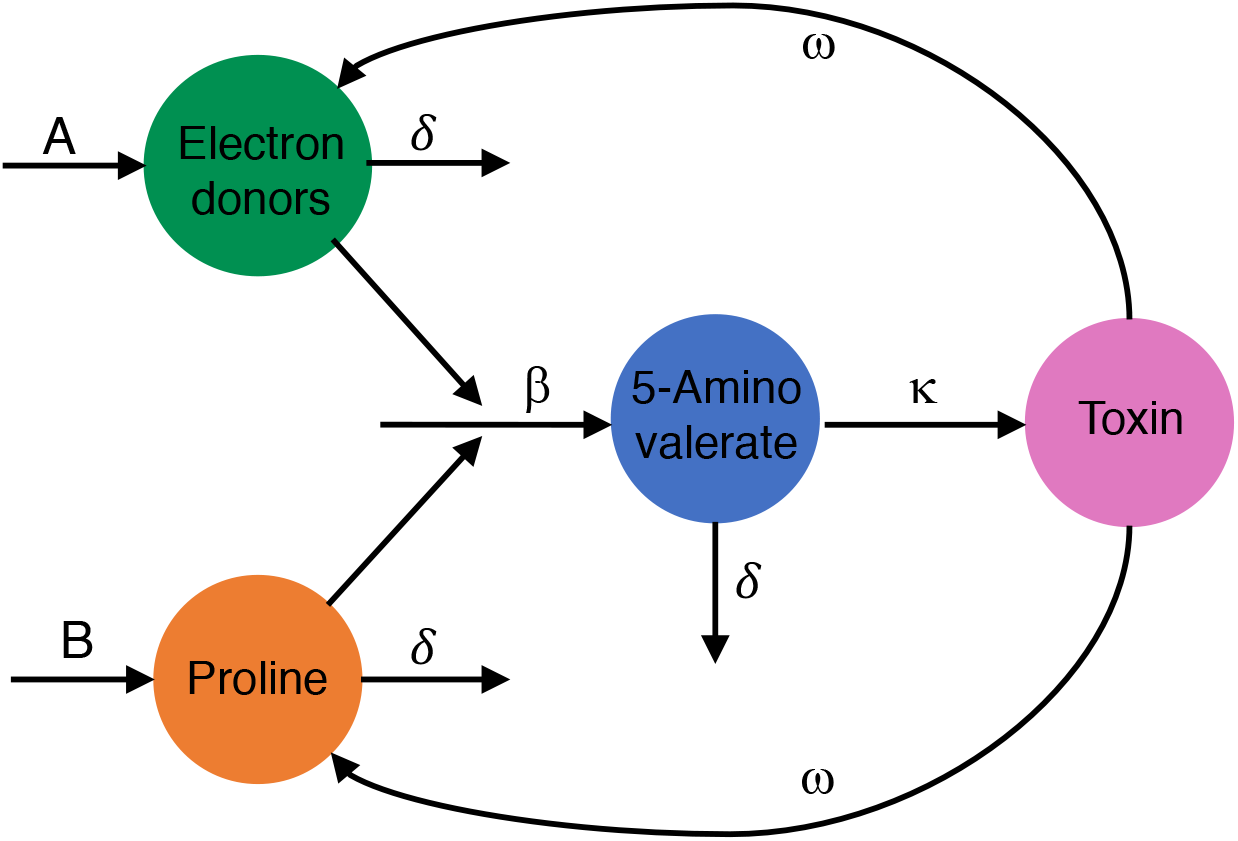
Schematic of the Stickland reaction that leads to *C. difficile* growth and subsequent toxin production.

These results highlight the need to further investigate biochemical mechanisms underpinning these relationships. Both proline and 5-aminovalerate are involved in the Stickland reaction which provides energy to *C. difficile* [42, 33, 43, 44]. Through this reaction, *C. difficile* proline reductase reduces proline to 5-aminovalerate by using an electron donated to NADH by an amino acid (typically branched chain amino acids) from the oxidative branch of the Stick-land reaction [52]. As proline levels decrease, the NAD+ generated by this reaction decreases, leading to a dramatic shift in *C. difficile* metabolism through the transcriptional regulator Rex, and an increase in toxin gene expression [53]. The other three metabolites connected to both *tcdA* and *tcdB* are sedoheptulose, guanosine, and hydroxyphenyl propionate. The negative association between sedoheptulose and toxin gene expression may indicate that the inflammation elicited by toxin activity is altering the immunometabolism of the gut tissue by polarizing macrophages to an M2-like phenotype via the activity of sedoheptulose kinase, thus leading to decreased sedoheptulose levels when toxin gene expression is high [54]. It is unclear at this time why guanosine and hydroxyphenyl propionate are negatively correlated with the expression of the toxin genes. The latter is a metabolite of microbial amino acid metabolism, so it is possible that the toxin-induced inflammation leads to an decrease in the level of the species that produce it.

### 3.2 ODE Results

A system of ODEs based on the results of the graphical analysis was developed to model the Stickland reaction, to further investigate its role in *C. difficile* infection and toxin production. The Stickland reaction involves the coupled deamination of two amino acids and is the principal chemical reaction by which *Clostridia* and other anaerobic bacteria obtain energy [52]. In the Stickland reaction, a hydrogen is transferred to NAD+ to produce NADH, which then supplies the reducing power to produce a Stickland product.

Graphical analysis suggests that the conversion of proline to 5-aminovalerate is temporally connected to *C. difficile* toxin gene expression (Figure 3). There is no direct indication of precisely which electron donor may be involved in this reaction. However, a broader approach was implemented to model general electron donors in our Stickland reaction model by considering the electron acceptor proline and its Stickland byproduct 5-aminovalerate as state variables (Figure 4). An ODE model was then developed from this schematic.

We developed an ODE model to better understand the components involved in the Stickland reaction by considering three primary metabolites: the electron donors as a group, proline, 5-aminovalerate; we also use toxin production for *C. difficile* growth. In the ODE model, the electron donors and proline are assumed to each have the same constant source term, *A* (metabolite h^*−*1^). All three populations—electron donors, proline, and 5-aminovalerate—are removed or eliminated at the same rate, *δ* (h^*−*1^), under the assumption of a well mixed environment. The electron donors and proline interact 1:1 at rate, *β* (metabolite^*−*1^h^*−*1^), using the proportion of electron donors to the total metabolites, instead of concentration quantities. It is also assumed that 5-aminovalerate leads to *C. difficile* growth and subsequent toxin production. Explicit bacterial growth of *C. difficile* is excluded from this model. This reduces the parameters space to enable more precise parameter estimation in the toxin feedback loop. While incorporating *C. difficle* growth explicitly could be beneficial, only the toxin expression in our model due to the limited data availability of that data.

The Stickland reaction that produces 5-aminovalerate, *S*, creates energy for *C. difficile* colonization. We assume *C. difficile* has a logistic growth rate, as previous bacterial models have used [55]. However since we only model toxin production, we include the logistic growth of bacteria in the toxin term and assume toxin production also has a logistic growth rate. The carrying capacity of toxin production is dependent on the total amount of 5-aminovalerate, (*KS*). Toxin is utilized to cause gut inflammation and destruction to the host at rate *ωT* (h^*−*1^), respectively. The model describing these interactions is given by the system:

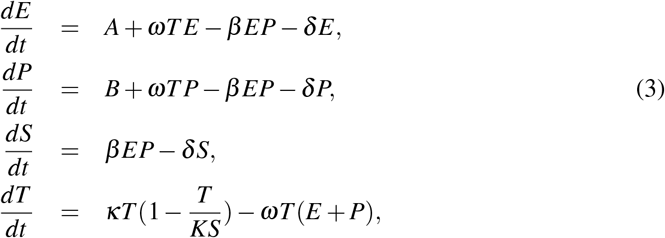

with initial conditions *E*(0) = *E*_0_ metabolites, *P*(0) = *P*_0_ metabolites, *S*(0) = *S*_0_ metabolites, and *T* (0) = *T*_0_ toxin. For numerical stability, a normalized derivation of this model was implemented and the specific details of this derivation can be found in Appendix A.

#### 3.2.1 Concentration data and model parameterization

The most efficient amino acid electron donors for the Stickland reactor are leucine, isoleucine, and valine [46]. It is assumed that the primary Stickland acceptor is proline and the primary Stickland product is 5-aminovalerate, based on the graphical analysis 3. Thus our *E* population is given by the sum of the normalized concentrations of leucine, isoleucine, and valine. *P* and *S* represent the normalized concentration of proline and 5-aminovalerate, respectively. The missing values in the raw ion data without pre-processing was first filled using the K-nearest neighbor algorithm prior to data normalization. Equation 4 was normalized so that each *E, P*, and *S* were normalized to the total *E* + *P* + *S*, explicitly: *Ê* = *E/*(*E* + *S* + *P*), 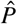 and *Ŝ* = *S/*(*E* + *S* + *P*). The toxin data was scaled so that *T* = ([*tcdA*] + [*tcdB*])*/*[*RpoC*]. The median values of the data for *Ê*, 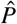, and *Ŝ* across all samples at *t* = 0 are used at the initial conditions for *Ê*_0_, 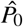, and *Ŝ*_0_. Finally, an MCMC-based parameter estimation was then performed to fit the normalized data to the normalized model using 100 walkers, each generating 1,000 samples each and utilizing a 100-sample burn-in (100,000 total samples).

#### 3.2.2 Numerical Results

The initial conditions for the electron, proline, and 5-aminovalerate, *Ê*_0_, 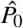, and *Ŝ*_0_, were selected to be the median value from the omics data (i.e. med(*Ê*(0)). To understand the sensitivity of the final amount of toxin to the parameters *T* at 30 h, *T* (30 h), we perform a global sensitivity analysis. This is done with Latin hypercube sampling and partial rank correlation coefficients (PRCC) [37]. A total of 100,000 simulations are executed. Additional information and figures are available in the supplemental material. Based on the PRCC analysis, the decay rate of all metabolites, *δ*, was identified as the parameter that affects the toxin amount at 30 h the least, and was therefore fixed to *δ* = 0.1 h^*−*1^. The toxin growth rate, *κ*, was also fixed since K and *κ* have indistinguishable effects and relatively little effect on *T* (30 h) based on the sensitivity analysis to *κ* = 1 h^*−*1^. The initial conditions and fixed parameters are summarized in Table 2.

**Table 2:** Fixed parameters used in the normalized ode system. The initial values were estimated from the concentration data.

| Parameter | Units | Value | Description |
| --- | --- | --- | --- |
| $\hat{E}_0$ | - | 0.432 | Initial concentration of electron donors |
| $\hat{P}_0$ | - | 0.563 | Initial concentration of proline |
| $\hat{S}_0$ | - | 0.004 | Initial concentration of 5-aminovalerate |
| $T_0$ | - | 0.001 | Initial amount of toxin |
| $\delta$ | $\text{h}^{-1}$ | 0.01 | Metabolite decay rate |
| $\kappa$ | $\text{h}^{-1}$ | 1 | Toxin production |

After estimating *A, B, K, β*, and *ω* for Equations (7), the Stickland reaction was numerically simulated to examine how each changed the normalized toxin level *T* at 30 h using MCMC methods. During this analysis, the posterior probability distribution of these parameters was sampled 100,000 total times (Figure 5). The 2-dimensional scatter plots of these samples are shown and demonstrate secondary dependencies between the fit parameters. These data show that each of the parameters are normally distributed and the mean and standard deviation of each parameter fit from the MCMC simulations are plotted in Figure 6 and summarized in Table 4. The relatively large standard errors associated with the parameter fits are likely due to the relatively few time-series measurements available as transcriptomic and metabolomic data was only collected at 0, 12, 24, and 30 h. Further, the data itself has a relatively high standard deviation between specimens, at each timepoint. Though the standard errors associated with the data fits for *β, A, B*, and *K*, they are less than the expected values obtaind from the regression. However, the largest relative uncertainty is associated with *ω*, which governs the feedback rate (Figure 4) and is one of the more sensitive parameters affecting the model conditions after 30 h (Figure A8). More experimental data is needed to reduce the uncertainty associated with *ω*, but the mechanistic model presented here has identified it as a critical parameter for follow-up studies to target.

**Table 3:**
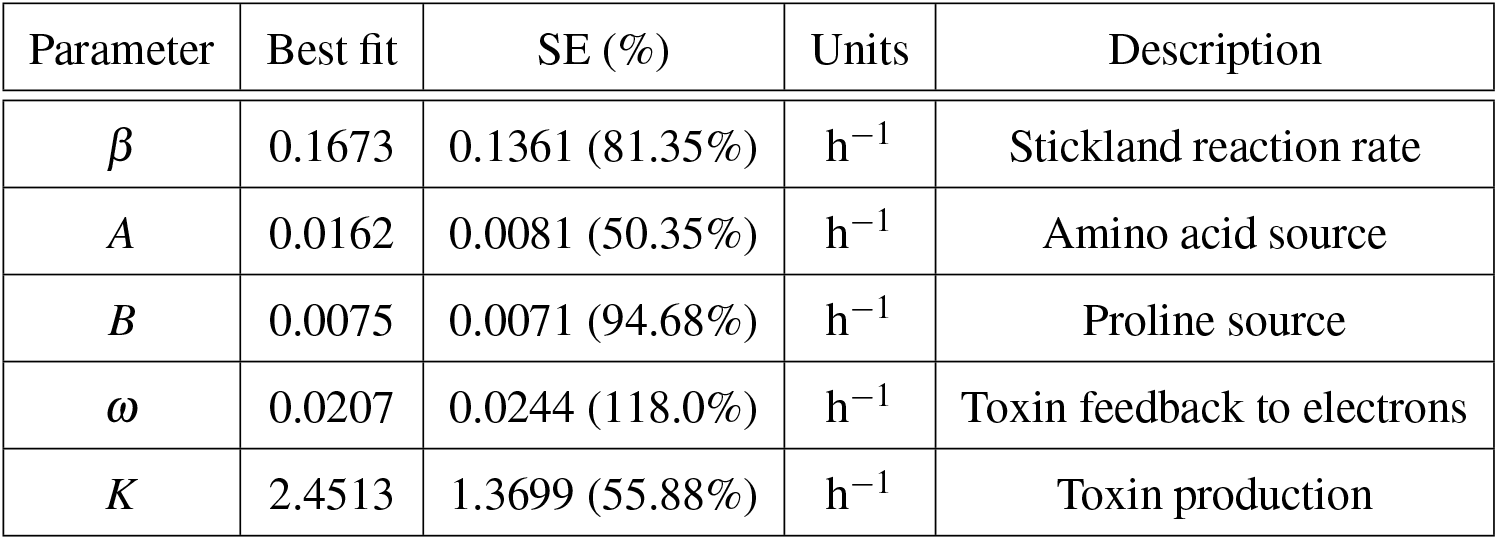
Best fit parameters described by the MCMC chain after 100,000 steps.

| Parameter | Best fit | SE (%) | Units | Description |
| --- | --- | --- | --- | --- |
| $\beta$ | 0.1673 | 0.1361 (81.35%) | $\text{h}^{-1}$ | Stickland reaction rate |
| $A$ | 0.0162 | 0.0081 (50.35%) | $\text{h}^{-1}$ | Amino acid source |
| $B$ | 0.0075 | 0.0071 (94.68%) | $\text{h}^{-1}$ | Proline source |
| $\omega$ | 0.0207 | 0.0244 (118.0%) | $\text{h}^{-1}$ | Toxin feedback to electrons |
| $K$ | 2.4513 | 1.3699 (55.88%) | $\text{h}^{-1}$ | Toxin production |

**Table 4:** Parameters, ranges, and baseline value for the sensitivity and uncertainty analysis.

| Parameter | Range | Baseline |
| --- | --- | --- |
| $A$ | $(10^{-3}, 1)$ | 0.1 |
| $B$ | $(10^{-3}, 1)$ | 0.1 |
| $\omega$ | $(10^{-3}, 1)$ | 0.1 |
| $\beta$ | $(10^{-3}, 1)$ | 0.1 |
| $\delta$ | $(10^{-3}, 1)$ | 1 |
| $K$ | $(10^{-2}, 100)$ | 1 |
| $\kappa$ | $(10^{-2}, 5)$ | 1 |

**Figure 5.**
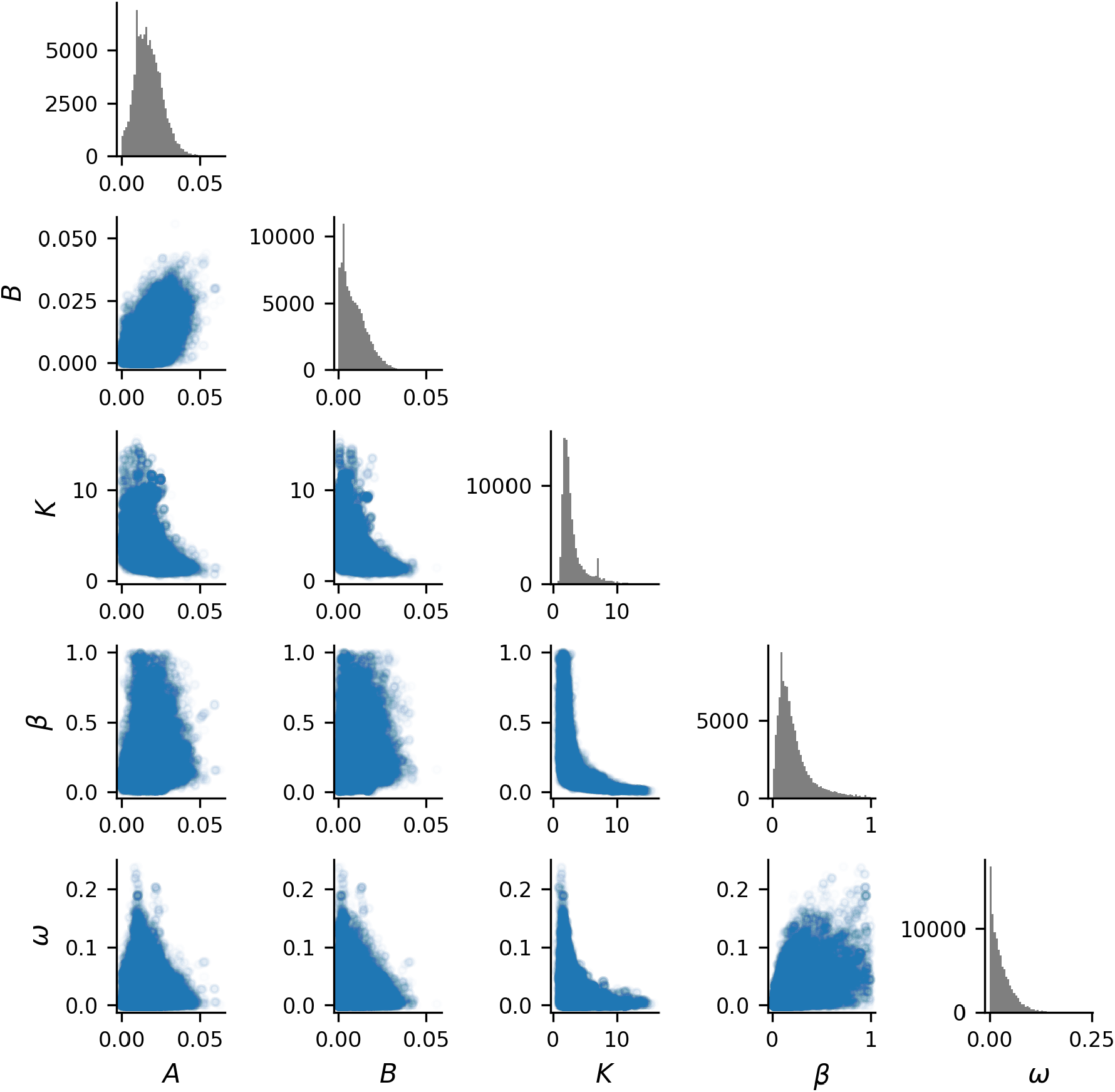
MCMC parameter estimation using 100,000 simulations. The histogram on the top of each column (gray) represents the distribution of simulated parameters around which the MCMC algorithm converged for the parameter labeled on the bottom of each column. The scatter plots (blue) represent the 2-dimensional distribution of each combination of parameter, labelled on the bottom and left axes.

**Figure 6.**
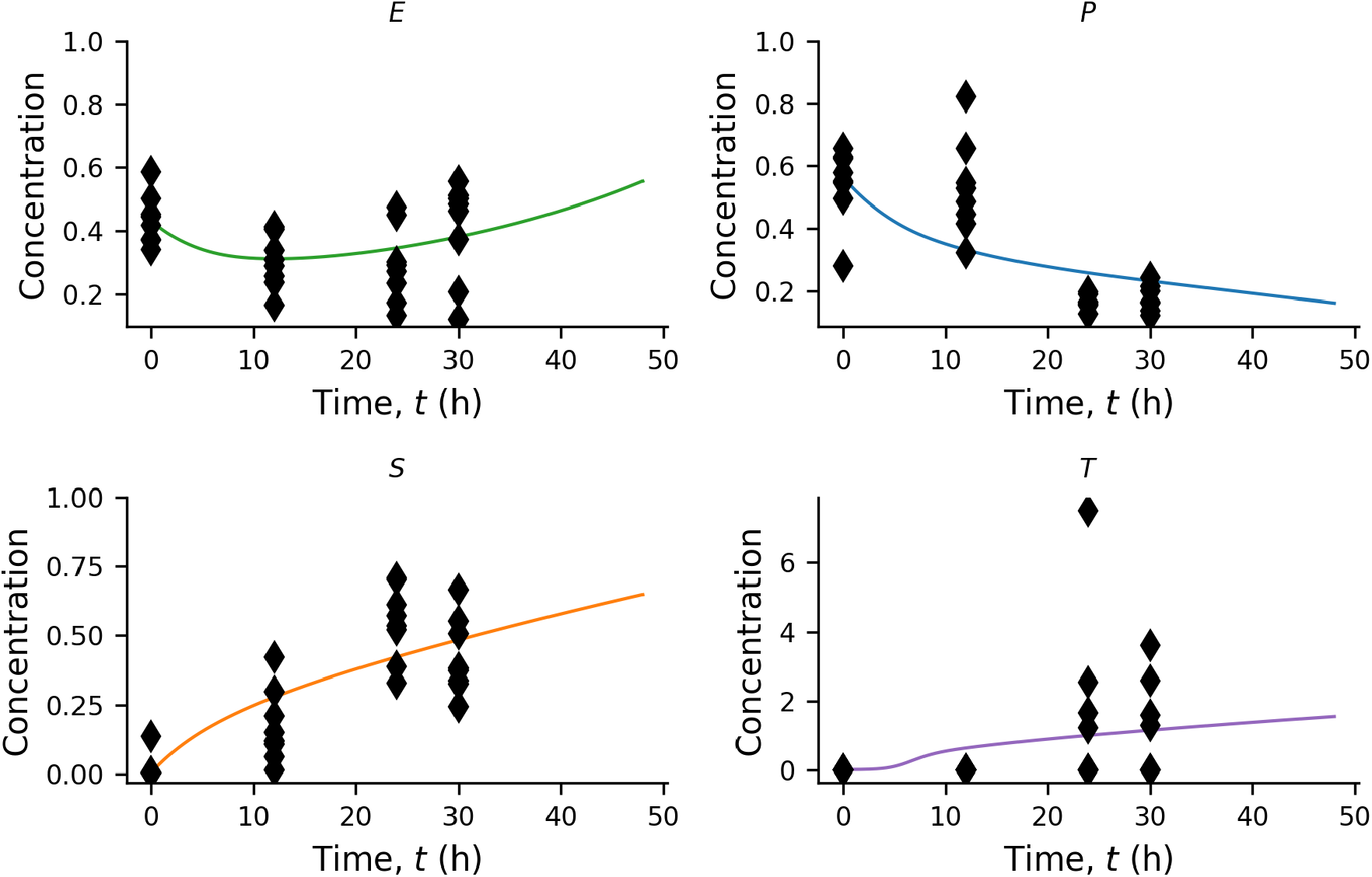
Solutions of model (7) using the parameters from our MCMC parameter estimation. We plot the electron donors, *E*, proline, *P*, 5-aminovalerate, *S*, and toxin, *T* . The relevant metabolomic data and toxin expression data are the diamonds (*◊*). The predictions are given in lines where the electron donors (green), proline (blue), 5-aminovalerate (orange) and toxin (purple).

Analyzing the fit of Equations 7 demonstrates that the best-fit parameterization is in good agreement with the experimental results (Figure 6. The electron donors, *E*, have a concentration of 0.39 after 30 hours, which is 0.04 lower than the proportion initially in the system. This suggests that the electron donor concentration is slightly disturbed by *C. difficile* colonization. Proline has a 30-h concentration of 0.28: half of the initial concentration. 5-aminovalerate concentration increases from a low initial concentration to a concentration of 0.48 of the metabolites after 30 h. This increases the concentration of Stickland products *S* with in the first hour of simulation, which provides energy for *C. difficile* and subsequent toxin production. The ODE model predicts that within the first 12 h there is a nominal increase in toxin, and at hours 24 and 30 or model has concentrations similar to that of our experimental data.

### 3.3 Analysis and interpretation

To identify the effects of modulating each of the fit parameters, the ODE model was solved for all parameters set to their best-fit values except for a single target parameter which was individually varied from 0.001-fold to 100-fold it best-fit value. Namely, *A, B, K, β, ω*, and *κ* were varied individually and the results of increasing and decreasing each parameter are shown in Figure 7. In each case, In the simulations 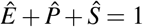 over all time, and no restriction is imposed on the toxin output and can thus grow very large. This parameter variation analysis demonstrates multiple targets for reducing toxin production at the 30 h time point, or when the study ends. Our overall goal is to decrease the final amount of toxin gene expression, and highlight potential mechanisms that could mitigate disease. The most noteworthy and testable finding from this analysis is that increases or decreases in *A* and *B* (top left and center panels in Figure 7), the amino acid source terms, lead to decreases in toxin. This suggest that disrupting the hosts amino acid make-up, specific to electron donors or proline, could be sufficient to limit toxin production. Decreasing *β* which would indicate slowing down the Stickland reaction would also reduce toxin. Additionally, increasing the feedback parameter *ω* leads to more usage of the toxin which ultimately leads to a decrease in the final amounts of toxin gene expression. Increasing *K* leads to an increased carrying capacity of toxin to a threshold. Because this term is modeled as a carrying capacity, numerically we see an instability of toxin once the carrying capacity overpowers the toxin term. Similar numerical instabilities are noted with variations of *κ* because this term is part of the same term in the 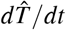 equation. Varying the parameters in the 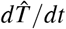 equation, *K* and *κ* are trivial cases, however this does indicate the model and relative parameter analysis is working as expected.

**Figure 7.**
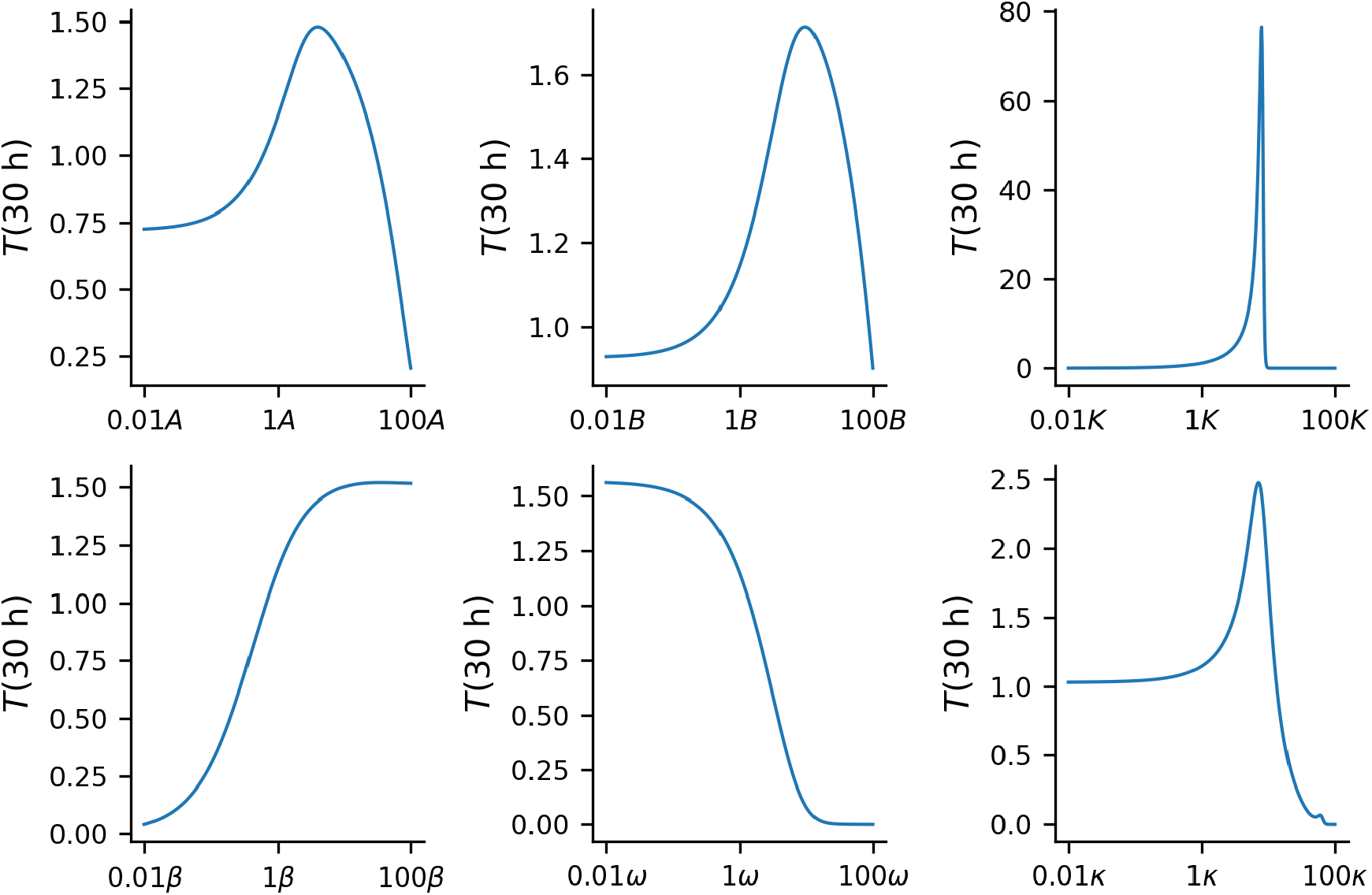
Individually adjusting individual parameters by two fold from the estimate (above and below) and predicting the change in toxin. The y-axis represents the amount of toxin after 30 hours, and the x-axis is the fold change from the estimated parameter vales

**Figure 8.**
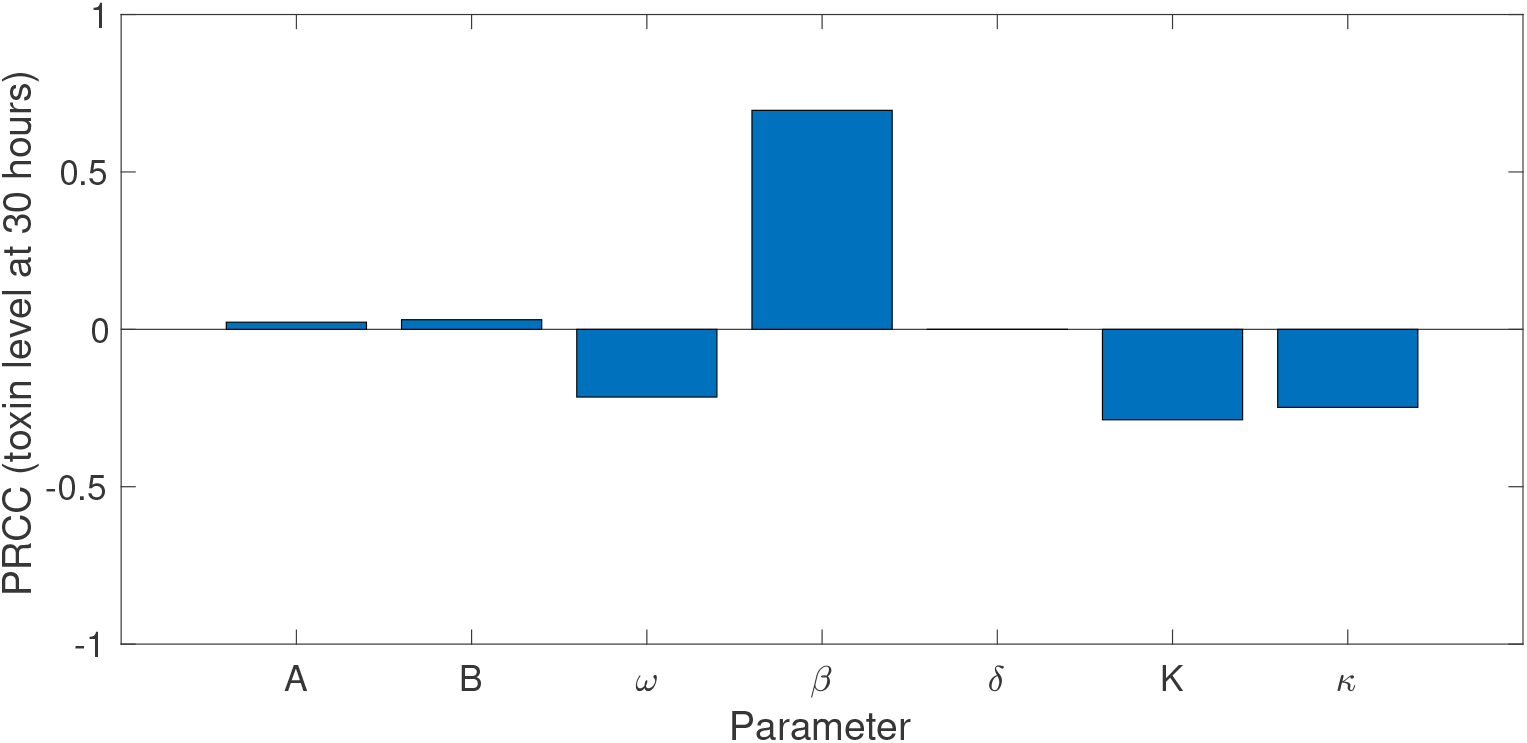
Sensitivity and uncertainty analysis using Latin hypercube sampling (LHS) of parameter space and partial rank correlation coefficients (PRCC). PRCC of total amount of toxin at 30 hours. All parameter ranges are provided in the supplemental table. A total of 100,000 simulations were executed to obtain all PRCC values.

## 4 Discussion

In this work, a novel pipeline for utilizing metabolomic data to drive the formulation of a mechanistic mathematical model was been implemented. First, the incomplete raw data was addressed by filling in the missing data using the K-nearest neighbors algorithm. Second, sparsification techniques were used to elucidate the strongest partial correlations within the raw data to highlight potential mechanisms driving toxin production during a *C. difficile* infection. Third, a mechanistic model was derived using a system of ordinary differential equations overlaid on the results from the graphical analysis. Parameter estimation techniques were used to identified parameters and fit the ODE model to metabolomic and transcriptomic data. Finally, sensitivity and parameter variation analyses were used to identify potential therapeutic targets; in this case, the reduction of the *tcdA* and *tcdB* gene transcripts 30 h post-infection. Through this pipeline, proline and 5-aminovalerate were identified as the strongest partially-correlated metabolites with *tcdA* and *tcdB* gene expression. This analysis identified that by varying the levels of proline, toxin gene expression can be increased or decreased, consistent with *in vitro* results [46, 56, 57, 58]. It was also found that the proline to 5-aminovalerate Stickland reaction is highly partially correlated with *C. difficile* infection. This finding is corroborated through additional experimental studies [42, 13, 59, 49]. The consistency of our results with these findings suggests that the analysis pipeline developed herein is a viable strategy for identifying biological mechanisms from multi-omics data are sufficiently robust.

While the present work has provided a novel pipeline analysis for generating mechanistic models from high dimensional data, there remain several limitations. Despite the high-dimensionality of the metabolite and transcriptomic data extracted from each sample, the small sample size of the metabolomic specimens (*n* = 8) limits the reliability of the glasso, as is apparent in the use of a large penalization and selection of *λ* = 0.8 was heuristic and based on the overall graph structure. In the present example, the primary goal of this analysis was to identify key nutrients involved in toxin expression, and as such, focused on selecting the most consistent edges associated with tcdA and tcdB. A more stable penalization could be identified in the future, though this is not possible with the small sample size in the present study.

Additionally, the choice of the Stickland reaction and the final construction of the mechanistic model were based on additional biological information, outside the multi-omics data . Ideally, in future utilization of omics data, a direct link between the glasso and a mechanistic model could automated and based solely on the sparse graph. There were also many simplifications made in the ordinary differential equation system, including removing the actual measurements of *C. difficile* growth. While the inclusion of this data could improve the model, this work is a formative first step toward the utilization of multi-omics data for mechanistic model development using sparsification techniques.

## 5 Conclusion

In this study, a methodological pipeline has been implemented to develop a mechanistic model from high-dimensional muilti-omics data has been implemented and analyzed. This pipeline was demonstrated using example metabolomic and transcriptomic data from an *in vivo* murine model of *C. difficile* infection to identify potential therapeutic targets. While several limitations were identified, the approach to data utilization and model development presented in this work proved useful and accurate, with the aim of being stepping stone for future model development efforts based on metabolomic data. The utility of this new methodology is highlight by the testable predictions the analysis of the ODE model generated in this work reveals and their agreement with other recent experimental results.

## A Model Normalization

We consider normalized populations such that:

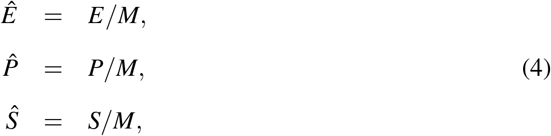

where *M* = *E* + *P* + *S*, then take the derivative of each normalized population with respect to *t* such that,

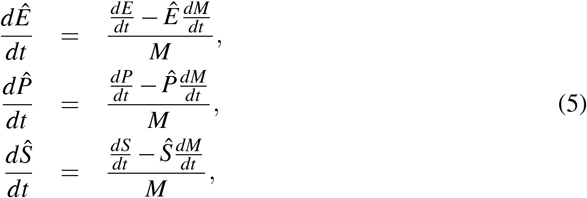

The change in the total population is given by the total change in the metabolites,

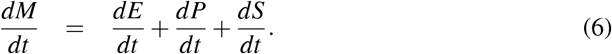

We now rewrite the original ODE model in terms of *Ê*,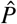, and Ŝ.

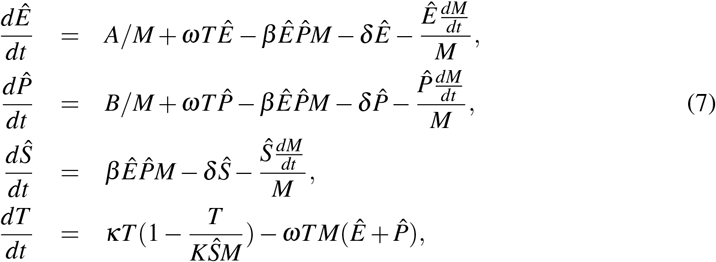

where,

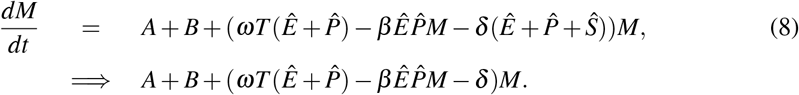

We assume a constant amount of metabolites within the gut which is corroborated by our experimental data, which stay in the same order of magnitude for the ion counts throughout the duration of our mathematical simulation.

The metabolite populations (*Ê*, 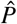, *Ŝ*) are now expressed as normalized concentrations. The toxin production is in terms of toxin per hour, albeit *tcdA* and *tcdB* is normalized by the *rpoC* housekeeping gene. Additionally this system maintains the original system dynamics from our biologically based model of the Stickland reaction.

## Notes

### Competing Interest Statement

The authors have declared no competing interest.

## References

[1] Adam M Feist and Bernhard Ø Palsson. The growing scope of applications of genome-scale metabolic reconstructions using escherichia coli. Nature biotechnology, 26(6):659–667, 2008.

[2] Matthew J Silk, Darren P Croft, Richard J Delahay, David J Hodgson, Nicola Weber, Mike Boots, and Robbie A McDonald. The application of statistical network models in disease research. Methods in Ecology and Evolution, 8(9):1026–1041, 2017.

[3] Jan Krumsiek, Karsten Suhre, Anne M Evans, Matthew W Mitchell, Robert P Mohney, Michael V Milburn, Brigitte Wägele, Werner Römisch-Margl, Thomas Illig, Jerzy Adamski, et al. Mining the unknown: a systems approach to metabolite identification combining genetic and metabolic information. 2012.

[4] SM Ciupe and JM Heffernan. In-host modeling. Infect Dis Model, 2:188–202, 2017.

[5] P Baccam, C Beauchemin, CA Macken, FG Hayden, and AS Perelson. Kinetics of influenza a virus infection in humans. J virol, 80:7590–9, 2006.

[6] CAA Beauchemin and A Handel. A review of mathematical models of influenza a infections within a host or cell culture: lessons learned and challenges ahead. BMC public health, 11:S7, 2011.

[7] AM Smith, FR Adler, RM Ribeiro, RN Gutenkunst, and et al. Kinetics of coinfection with influenza a virus and streptococcus pneumoniae. PLoS path, 9:e1003238, 2013.

[8] MA Stafford, L Corey, Y Cao, ES Daar, DD Ho, and AS Perelson. Modeling plasma virus concentration during primary hiv infection. J theor biol, 203:285–301, 2000.

[9] Samantha Erwin and Stanca M Ciupe. Germinal center dynamics during acute and chronic infection. Mathematical Biosciences & Engineering, 14(3):655, 2017.

[10] J Forde S Ciupe, A Cintron-Arias, and S Lenhart. Optimal control of drug therapy in a hepatitis b model. Appl Sci, 6:219, 2016.

[11] Samantha Erwin, Derek M Foster, Megan E Jacob, Mark G Papich, and Cristina Lanzas. The effect of enrofloxacin on enteric escherichia coli: Fitting a mathematical model to in vivo data. PLoS One, 15(1):e0228138, 2020.

[12] E Herrmann, AU Neumann, JM Schmidt, and S Zeuzem. Hepatitis c virus kinetics. Antivir ther, 5:85–90, 2000.

[13] JR Fletcher, S Erwin, C Lanzas, and CM Theriot. Shifts in the Gut Metabolome and Clostridium difficile Transcriptome throughout Colonization and Infection in a Mouse Model. mSphere, 3:1–18, 2018.

[14] KT Do, i Kastenmller, DO Mook-Kanamori, NA Yousri, and et al. Network-based approach for analyzing intra-and interfluid metabolite associations in human blood, urine, and saliva. J of Proteome Res, 14(2):1183–1194, 2015.

[15] H Chen, SA Quandt, JG Grzywacz, and TA Arcury. A distribution-based multiple imputation method for handling bivariate pesticide data with values below the limit of detection. Environ health perspectives, 119:351–356, 2011.

[16] J Xia, N Psychogios, N Young, and DS Wishart. Metaboanalyst: a web server for metabolomic data analysis and interpretation. Nucleic acids res, 37:W652–W660, 2009.

[17] Michael X Chen, San-Yuan Wang, Ching-Hua Kuo, and I-Lin Tsai. Metabolome analysis for investigating host-gut microbiota interactions. Journal of the Formosan Medical Association, 118:S10–S22, 2019.

[18] KT Do, S Wahl, J Raffler, S Molnos, and et al. Characterization of missing values in untargeted ms-based metabolomics data and evaluation of missing data handling strategies. Metabolomics, 14:128, 2018.

[19] Javier E Flores, Daniel M Claborne, Zachary D Weller, Bobbie-Jo M Webb-Robertson, Katrina M Waters, and Lisa M Bramer. Missing data in multi-omics integration: Recent advances through artificial intelligence. Frontiers in Artificial Intelligence, 6, 2023.

[20] J Fan, F Han, and H Liu. Challenges of big data analysis. Natl sci rev, 1(2):293–314, 2014.

[21] CM Carvalho, J Chang, JE Lucas, JR Nevins, Q Wang, and M West. High-dimensional sparse factor modeling: applications in gene expression genomics. J Am Stat Assoc, 103:1438–56, 2008.

[22] H Rue and L Held. Gaussian Markov random fields: theory and applications. CRC press, 2005.

[23] A Dobra, C Hans, B Jones, JR Nevins, G Yao, and M West. Sparse graphical models for exploring gene expression data. J Multivar Anal, 90:196–212, 2004.

[24] F Rohart, B Gautier, A Singh, and KA Le Cao. mixomics: An r package for omics feature selection and multiple data integration. PLoS Comput Biol, 13:e1005752, 2017.

[25] Y Zhu and I Cribben. Sparse graphical models for functional connectivity networks: Best methods and the autocorrelation issue. Brain connect, 8:139–65, 2018.

[26] R Tibshirani. Regression shrinkage and selection via the lasso. J Royal Statist Soc B, 58:267–88, 1996.

[27] J Friedman, T Hastie, and R Tibshirani. Sparse inverse covariance estimation with the graphical lasso. Biostatistics, 9:432–41, 2008.

[28] H Liu, F Han, M Yuan, J Lafferty, and L Wasserman. High-dimensional semiparametric gaussian copula graphical models. Annals Stats, 40:2293–326, 2012.

[29] C Gabor and N Tamas. The igraph software package for complex network research. Inter Journal, Complex Systems:1695, 2006.

[30] Franz-Georg Wieland, Adrian L Hauber, Marcus Rosenblatt, Christian Tönsing, and Jens Timmer. On structural and practical identifiability. Current Opinion in Systems Biology, 25:60–69, 2021.

[31] Joshua R Fletcher, Colleen M Pike, Ruth J Parsons, Alissa J Rivera, Matthew H Foley, Michael R McLaren, Stephanie A Montgomery, and Casey M Theriot. Clostridioides difficile exploits toxin-mediated inflammation to alter the host nutritional landscape and exclude competitors from the gut microbiota. Nature communications, 12(1):462, 2021.

[32] Casey M Theriot, Joshua R Fletcher, et al. Human fecal metabolomic profiling could inform clostridioides difficile infection diagnosis and treatment. The Journal of Clinical Investigation, 129(9):3539–3541, 2019.

[33] AD Reed, JR Fletcher, YY Huang, R Thanissery, AJ Rivera, RJ Parsons, et al. The stick-land reaction precursor trans-4-hydroxy-l-proline differentially impacts the metabolism of clostridioides difficile and commensal clostridia. msphere. 2022; 7: e0092621.

[34] Samantha Erwin, Lauren M Childs, and Stanca M Ciupe. Mathematical model of broadly reactive plasma cell production. Scientific Reports, 10(1):1–12, 2020.

[35] AC Hindmarsh and LR Petzold. Lsoda, ordinary differential equation solver for stiff or non-stiff system. 2005.

[36] Jonathan Goodman and Jonathan Weare. Ensemble samplers with affine invariance. Communications in applied mathematics and computational science, 5(1):65–80, 2010.

[37] S Marino, IB Hogue, CJ Ray, and DE Kirschner. A methodology for performing global uncertainty and sensitivity analysis in systems biology. J Theor Biol, 254:178–96, 2008.

[38] Meghna Verma, Samantha Erwin, Vida Abedi, Raquel Hontecillas, Stefan Hoops, Andrew Leber, Josep Bassaganya-Riera, and Stanca M Ciupe. Modeling the mechanisms by which hiv-associated immunosuppression influences hpv persistence at the oral mucosa. PloS one, 12(1):e0168133, 2017.

[39] R Core Team. R: A Language and Environment for Statistical Computing. R Foundation for Statistical Computing, Vienna, Austria, 2017.

[40] RStudio Team. RStudio: Integrated Development Environment for R. RStudio, Inc., Boston, MA, 2019.

[41] Guido Van Rossum and Fred L Drake Jr. Python reference manual. Centrum voor Wiskunde en Informatica Amsterdam, 1995.

[42] ML. Jenior, JL. Leslie, VB. Young, and PD. Schloss. Clostridium difficile colonizes alternative nutrient niches during infection across distinct murine gut microbiomes. mSystems, 2, 2017.

[43] Andrea Martinez Aguirre, Nazli Yalcinkaya, Qinglong Wu, Alton Swennes, Mary Elizabeth Tessier, Paul Roberts, Fabio Miyajima, Tor Savidge, and Joseph A Sorg. Bile acid-independent protection against clostridioides difficile infection. PLoS Pathogens, 17(10):e1010015, 2021.

[44] Laura Cersosimo, Madeline Graham, Auriane Monestier, Aidan Pavao, Jay N Worley, Johann Peltier, Bruno Dupuy, and Lynn Bry. Central in vivo mechanisms by which c. difficile’s proline reductase drives efficient metabolism, growth, and toxin production. bioRxiv, pages 2023–05, 2023.

[45] Sarah Jackson, Mary Calos, Andrew Myers, and William T Self. Analysis of proline reduction in the nosocomial pathogen clostridium difficile. Journal of bacteriology, 188(24):8487–8495, 2006.

[46] L Bouillaut, WT Self, and AL Sonenshein. Proline-dependent regulation of Clostridium difficile stickland metabolism. J of bacteriol, 195:844–854, 2013.

[47] Henning Dannheim, Thomas Riedel, Meina Neumann-Schaal, Boyke Bunk, Isabel Schober, Cathrin Spröer, Cynthia Maria Chibani, Sabine Gronow, Heiko Liesegang, Jörg Overmann, et al. Manual curation and reannotation of the genomes of clostridium difficile 630δ erm and c. difficile 630. Journal of Medical Microbiology, 66(3):286–293, 2017.

[48] Meina Neumann-Schaal, Julia Danielle Hofmann, Sabine Eva Will, and Dietmar Schomburg. Time-resolved amino acid uptake of clostridium difficile 630δ erm and concomitant fermentation product and toxin formation. BMC microbiology, 15:1–12, 2015.

[49] Eric J Battaglioli, Vanessa L Hale, Jun Chen, Patricio Jeraldo, Coral Ruiz-Mojica, Bradley A Schmidt, Vayu M Rekdal, Lisa M Till, Lutfi Huq, Samuel A Smits, et al. Clostridioides difficile uses amino acids associated with gut microbial dysbiosis in a subset of patients with diarrhea. Science translational medicine, 10(464):eaam7019, 2018.

[50] Laurent Bouillaut, Thomas Dubois, Michael B Francis, Nadine Daou, Marc Monot, Joseph A Sorg, Abraham L Sonenshein, and Bruno Dupuy. Role of the global regulator rex in control of nad+-regeneration in clostridioides (clostridium) difficile. Molecular microbiology, 111(6):1671–1688, 2019.

[51] Michael A Johnstone and William T Self. d-proline reductase underlies proline-dependent growth of clostridioides difficile. Journal of Bacteriology, 204(8):e00229–22, 2022.

[52] B Nisman. The stickland reaction. Bacteriol rev, 18:16, 1954.

[53] Laurent Bouillaut, Thomas Dubois, Michael B Francis, Nadine Daou, Marc Monot, Joseph A Sorg, Abraham L Sonenshein, and Bruno Dupuy. Role of the global regulator rex in control of nad+-regeneration in clostridioides (clostridium) difficile. Molecular microbiology, 111(6):1671–1688, 2019.

[54] A Haschemi, P Kosma, L Gille, CR Evans, and et al. The sedoheptulose kinase carkl directs macrophage polarization through control of glucose metabolism. Cell metab, 15.

[55] AM Smith and AP Smith. A critical, nonlinear threshold dictates bacterial invasion and initial kinetics during influenza. Sci Rep, 6:38703, 2016.

[56] K Yamakawa, S Kamiya, X Meng, T Karasawa, and S Nakamura. Toxin production by Clostridium difficile in a defined medium with limited amino acids. J med microbiol, 41:319–23, 1994.

[57] D Ikeda, T Karasawa, K Yamakawa, R Tanaka, M Namiki, and S Nakamura. Effect of isoleucine on toxin production by Clostridium difficile in a defined medium. Zentralblatt Bakteriol, 287:375–86, 1998.

[58] S Karlsson, A Lindberg, E Norin, LG Burman, and T Åkerlund. Toxins, butyric acid, and other short-chain fatty acids are coordinately expressed and down-regulated by cysteine in Clostridium difficile. Infect and immun, 68(10):5881–8, 2000.

[59] CM Theriot, MJ Koenigsknecht, PE Carlson, GE Hatton, and et al. Antibiotic-induced shifts in the mouse gut microbiome and metabolome increase susceptibility to clostridium difficile infection. Nat commun, 5:1–10, 2014.

